# Enhanced binding of guanylated poly(A) RNA by the LaM domain of LARP1

**DOI:** 10.1101/2024.05.08.593157

**Authors:** Guennadi Kozlov, Jianning Jiang, Tyler Rutherford, Anne M. Noronha, Christopher J. Wilds, Kalle Gehring

## Abstract

La-related proteins (LARPs) are a family of RNA-binding proteins that share a conserved La motif (LaM) domain. LARP1 plays a role in regulating ribosomal protein synthesis and stabilizing mRNAs and has a unique structure without an RNA binding RRM domain adjoining the LaM domain. In this study, we investigated the physical basis for LARP1 specificity for poly(A) sequences and observed an unexpected bias for sequences with single guanines. Multiple guanine substitutions did not increase the affinity, demonstrating preferential recognition of singly guanylated sequences. We also observed that the cyclic di- nucleotides in the cCAS/STING pathway, cyclic-di-GMP and 3’,3’-cGAMP, bound with sub-micromolar affinity. Isothermal titration measurements were complemented by high-resolution crystal structures of the LARP1 LaM with six different RNA ligands, including two stereoisomers of a phosphorothioate linkage. The selectivity for singly substituted poly(A) sequences suggests LARP1 may play a role in the stabilizing effect of poly(A) tail guanylation.

## INTRODUCTION

La-related protein 1 (LARP1) is a member of the LARP family of RNA-binding proteins that includes LARP1B, LARP3 (genuine La protein), LARP4, LARP4B, LARP6, and LARP7. Each member has distinct functional roles in RNA metabolism and gene expression regulation [1, 2]. All LARPs share a highly conserved La motif (LaM) domain, which is generally accompanied by a downstream RNA recognition motif (RRM) [3, 4]. This tandem arrangement constitutes the La-module [5, 6]. Among human proteins, LARP1 and the related LARP1B are the only members that do not possess the RRM domain [7]. Instead, the region following the LaM domain contains a PAM2 motif (PABP-interacting motif 2), which binds poly(A)-binding protein C1 (PABPC1) [2]. The other well-characterized domain in LARP1 is the C-terminal DM15 domain, which binds the mRNA 5’ m7G cap and oligopyrimidine (TOP) motifs [2, 8]. Functionally, LARP1 regulates the stability, localization, and translation of TOP mRNAs downstream of mTORC1 signaling [9–12]. TOP mRNAs encode translation machinery components, such as ribosomal proteins and factors. LARP1 is directly phosphorylated by the mTOR kinase [13–17]. LARP1 also interacts with a large number of mRNAs in mTOR-independent pathways and has been implicated in cancer [18–23].

The LaM and RRM domains of LARP3, LARP4, LARP6, and LARP7 and the LaM domain of LARP1 have been structurally characterized [7, 24–33]. In LARP3, the LaM and RRM domains form a synergistic RNA-binding scaffold to bind uridylate oligonucleotides. The 3’-terminal uridylate binds to the LaM domain and the 3’ penultimate nucleotide interacts with a V-shaped clamp formed between the LaM and RRM domains [25, 26]. In contrast, the LARP1 LaM binds RNA as a stand-alone domain with a preference for poly(A) sequences [7]. The terminal and penultimate adenylate nucleotides (numbered -1 and -2 from the 3’-end) bind in the same way as uridylates in other LARP proteins but without contacts from an adjacent RRM domain. The terminal (-1) ribose is positioned by polar contacts with Asp346 and the adenine ring stacks on Phe348. The penultimate (-2) adenine ring stacks on Tyr336 and is positioned by a hydrogen bond with Gln333. Considerable variability was observed in the positions of the preceding nucleotides with either the (-3) or (-4) nucleotide stacking between His368 and the (-1) adenine.

LARP1 has been linked to protection of poly(A) from deadenylation and associated mRNA stabilization [2, 7, 34, 35]. Consistent with its association with PABP, LARP1 from human cell extracts immunoprecipitates with poly(A) RNA, but not with poly(U), poly(C), or poly(G) [36]. Assays in human HEK293 cells deleted of endogenous LARP1 (KO) showed that expression of LARP1 led to net lengthening of mRNA poly(A) tails which was attributed to protection from binding to the LaM domain [7, 37]. LARP1 in complex with PABPC1 was found preferentially associated with short tails and its depletion resulted in their accelerated deadenylation [35].

In the present study, we show that isolated guanosine residues in poly(A) sequences improve binding to the LaM domain of LARP1. High-resolution crystal structures of LARP1 LaM with six different RNAs: A_5_G, A_4_GA, A_3_GA_2_, U_6_, and (*R*_P_)- and (*S*_P_)-diastereomers of a phosphorothioate derivative reveal the molecular interactions that allow for plasticity in base recognition while maintaining specificity for the RNA 3’-end. The increased affinity of the LARP1 LaM domain for poly(A) sequences containing single guanine residues suggests that in vivo LARP1 should be enriched on guanylated mixed-tailed mRNAs.

## RESULTS

### NMR studies with single nucleotides expose binding preference for guanine

We carried out NMR titrations with single nucleotides binding to ^15^N-labeled LARP1 LaM to isolate the individual nucleotide interactions (Fig. 1A). The addition of nucleotides caused spectral changes similar to those observed with the hexamer A_6_ but with smaller shifts and fewer signals affected (Suppl. Fig. S1) [7]. Unlike A_6_, which showed NMR signals in slow exchange, the nucleotide titrations displayed fast exchange kinetics and significant broadening of some signals. The signals with the largest changes were the amide of Ile364 and side chain amides of Gln333.

**Figure 1.**
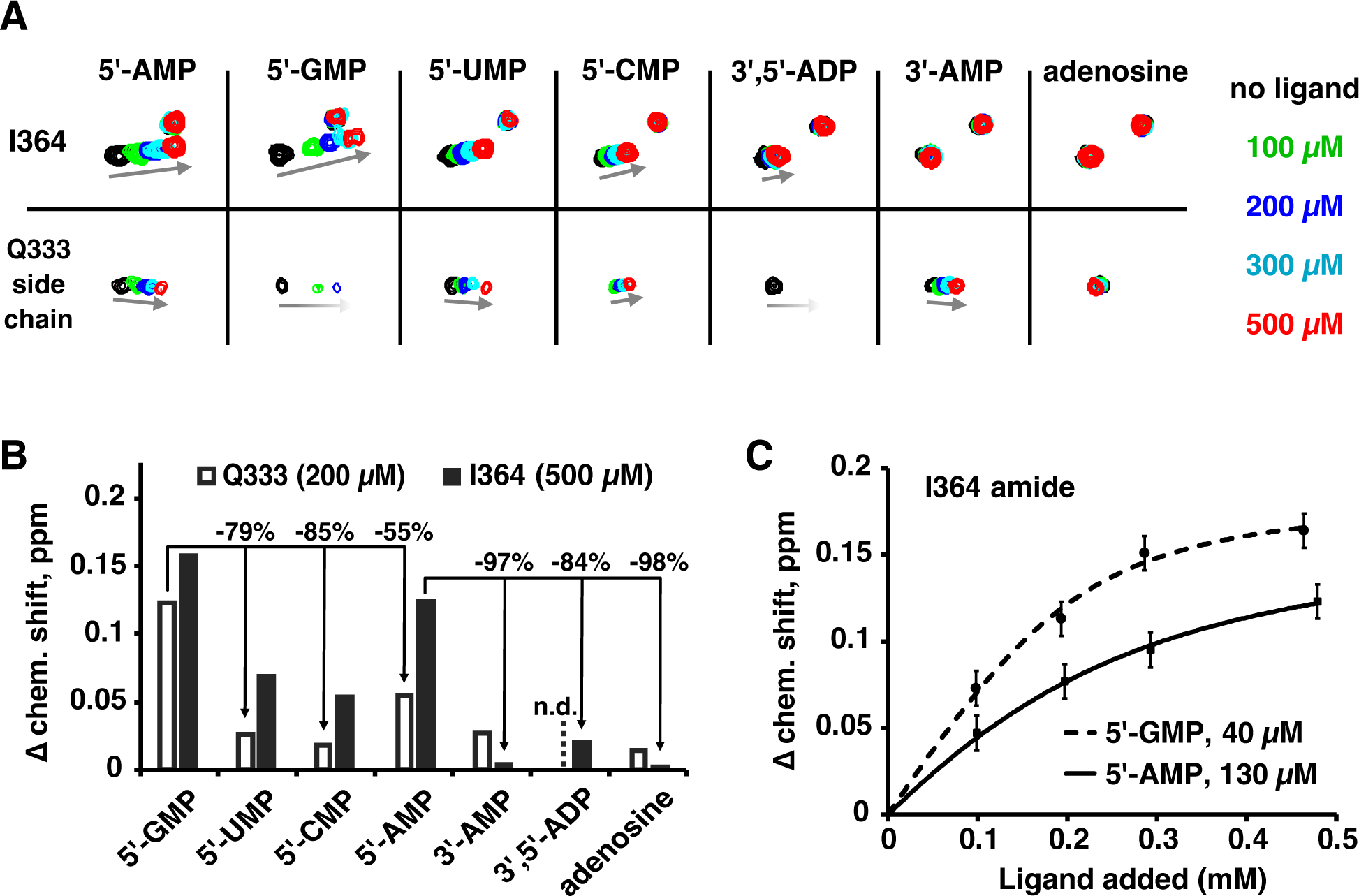
NMR reveals the LaM domain has distinct sites for mononucleotides with a preference for 5’-GMP. (A) Portions of ^15^N-^1^H correlation spectra of ^15^N-labeled LARP1 (323-410) upon addition of nucleotides or adenosine. Nucleotides bearing 3’-phosphates only shifted the Q333 side chain signal while 5’-phosphonucleotides shifted both Q333 and I364. Full spectra are shown in Suppl. Fig. S1. (**B**) Plot of the magnitude of proton chemical shift changes shows 5’-GMP induced the largest chemical shift changes. n.d.= not determined. (**C**) Measurement of binding affinities for 5’-GMP and 5’-AMP.

Unexpectedly, 5’-GMP showed larger chemical shift changes than 5’-AMP (Fig. 1B). At 200 µM, 5’-GMP shifted the Gln333 signal twice as much as 5’-AMP. Both purines nucleotides bound better (showed larger shifts) than the pyrimidines 5’-UMP and 5’-CMP. We used the size of the chemical shift change as a function of nucleotide concentration to estimate binding affinities (Fig. 1C). As the protein concentration is comparable to the binding affinity, fitting the changes required accounting for bound ligand when calculating the free ligand concentration. This had the advantage that it also provided an estimate of the binding stoichiometry. For 5’-GMP, the stoichiometry was measured to be 2.04 with an affinity of 40 µM. For 5’-AMP, the stoichiometry could not be determined but the *K*_d_ was measured to be 130 µM assuming a stoichiometry of 2. This is at odds with the affinities of oligonucleotides measured by isothermal titration calorimetry (ITC) [7]. The analysis of 22 different RNA oligonucleotides identified poly(A) RNA as the preferred ligand: A_6_ bound with 14-fold better affinity than G_6_, 11-fold better than U_6_, and 200-fold better than C_6_.

In addition to base specificity, the LaM domain is strongly selective for RNA with a 3’-hydroxyl. ITC experiments show that addition of a 3’-phosphate to A_6_ decreases the binding affinity 200-fold [7]. The NMR experiments showed good agreement: the shift of the Ile364 peak with 3’-AMP shift was 30-fold smaller than with 5’-AMP (Fig. 1B). Titrations with 3’,5’-ADP and adenosine confirmed the roles of the 3’-hydroxyl and 5’-phosphate, respectively.

Comparison of the shifts of Ile364 and Gln333 suggests that they reflect distinct nucleotide binding sites. For several ligands, the size of chemical shift changes for Ile364 and Gln333 differed markedly. Ile364 was not appreciably shifted by 3’-AMP while Gln333 was. Similarly, doubly charged 3’,5’-ADP induced a small shift in Ile364 but broadened the Gln333 signal beyond detection. In crystal structures, the Gln333 side chain makes a hydrogen bond with the base in the (-2) position which suggests the NMR shift primarily reflects occupancy at that position. The amide of Ile364 sits under Phe367 near the binding site of the (-1) nucleotide and likely reports on the occupancy of that site. This interpretation rationalizes the differences in ligand sensitivity. Ile364 is extremely sensitive to modification of the 3’-hydroxyl since it reflects binding of the 3’-terminal nucleotide, while Gln333 tolerates a 3’-phosphate. Conversely, Gln333 is involved in the extensive contacts with the (-2) nucleotide base and shows strong base selectivity, while Ile364 shows shifts of more equal size for all four nucleotide bases.

### Guanine improves RNA binding

We used isothermal titration calorimetry (ITC) experiments to probe the effect of guanine bases on RNA oligomer binding (Fig. 2, Suppl. Table S1, Suppl. Fig. S2). As previously reported, the LARP1 LaM domain bound the hexamer A_6_ with 250 nM affinity. Single guanine substitutions markedly improved binding with AAAAGA showing the highest affinity, 80 nM, a 3-fold improvement over A_6_. Substitutions at the (-1) and (-3) positions showed slightly smaller, approximately 2-fold increases. We also tested the effect of multiple guanine substitutions. Surprisingly, a second guanine base did not improve binding and, in some cases, decreased the affinity. For example, the doubly substituted hexamer AAAGAG bound with essentially the same affinity as AAAGAA and AAAAAG, while AAAAGG bound with half the affinity of AAAAGA.

**Figure 2.**
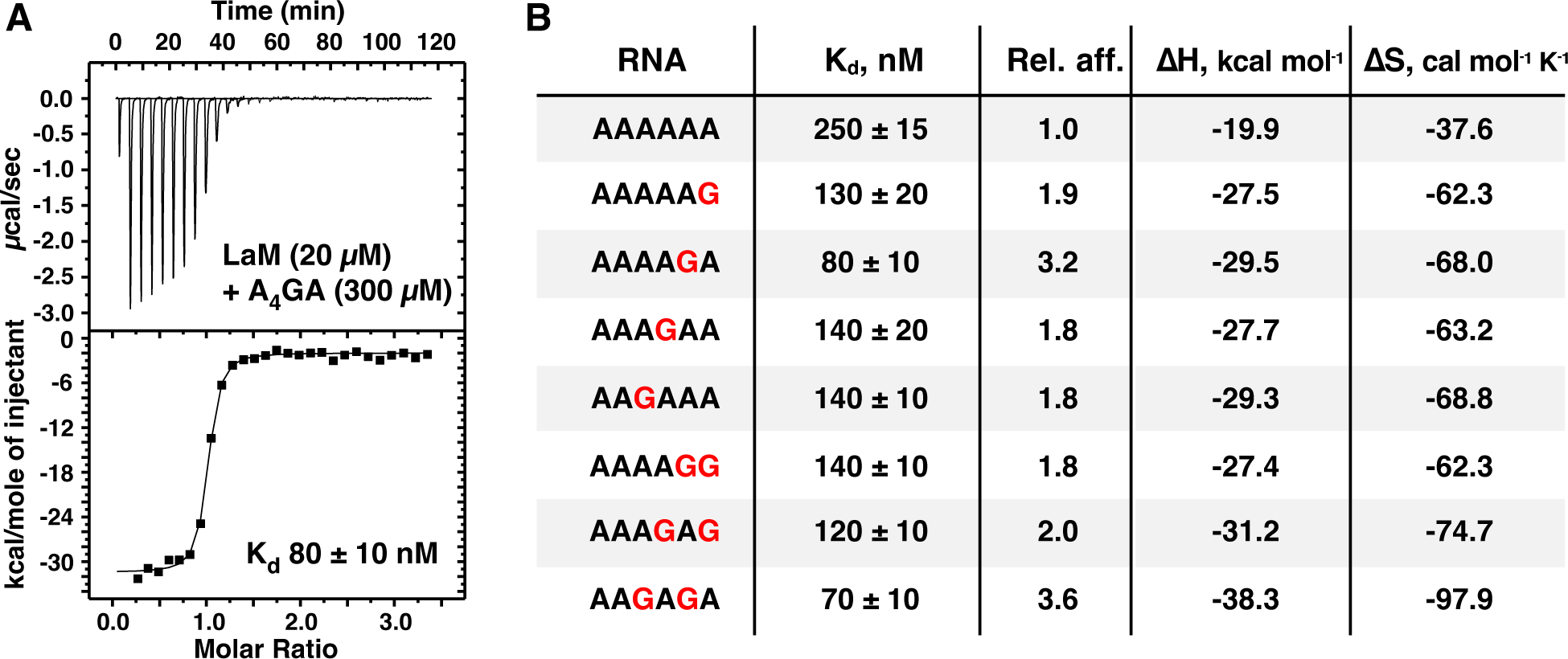
Isolated guanine bases increase the affinity of LaM binding measured by ITC. (A) Thermogram of the RNA ligand 5’-A_P_A_P_A_P_A_P_G_P_A-3’ binding LARP1 (323-410). **(B)** Table of binding affinities of singly and doubly guanine-substituted poly(A) RNAs. Experimental details are given in Suppl. Table 1 and Suppl. Fig. S2.

Previous ITC experiments measured an affinity of 3.3 µM for G_6_ binding to LaM, which is 40-fold weaker than AAAAGA. To explain the stark difference in affinity of single and all guanine RNAs, we examined the binding affinity of the dinucleotide GG by ITC. Injections of GG RNA into LaM generated a complex thermogram inconsistent with a single binding reaction (Fig. 3). Given the propensity of guanine oligonucleotides to form quadruplexes [38], we suspected that the complex thermogram arose from the presence of two processes: i) dissociation of quadruplexes of GG present in the concentrated solution in the syringe, and ii) binding of the single-stranded GG to the LaM domain. To test this, we reversed the concentrations of the components, injecting concentrated LaM into a 20-fold diluted solution of the RNA dinucleotide. This resulted in a normal thermogram which could be fit as a simple bimolecular interaction with 2.8 µM affinity. As a control, an ITC experiment with AG carried out in the standard manner measured an affinity of 3.1 µM, confirming the unusual titration behavior was specific to GG. As the stability of guanine quadruplexes increases with the number of guanines, these results explain the low affinity of G_6_ relative to A_6_ and the RNAs with only one or two guanine bases. Competition between quadruplex formation and LaM-binding decreases the apparent affinity of guanine-rich RNAs for the LARP1 LaM.

**Figure 3.**
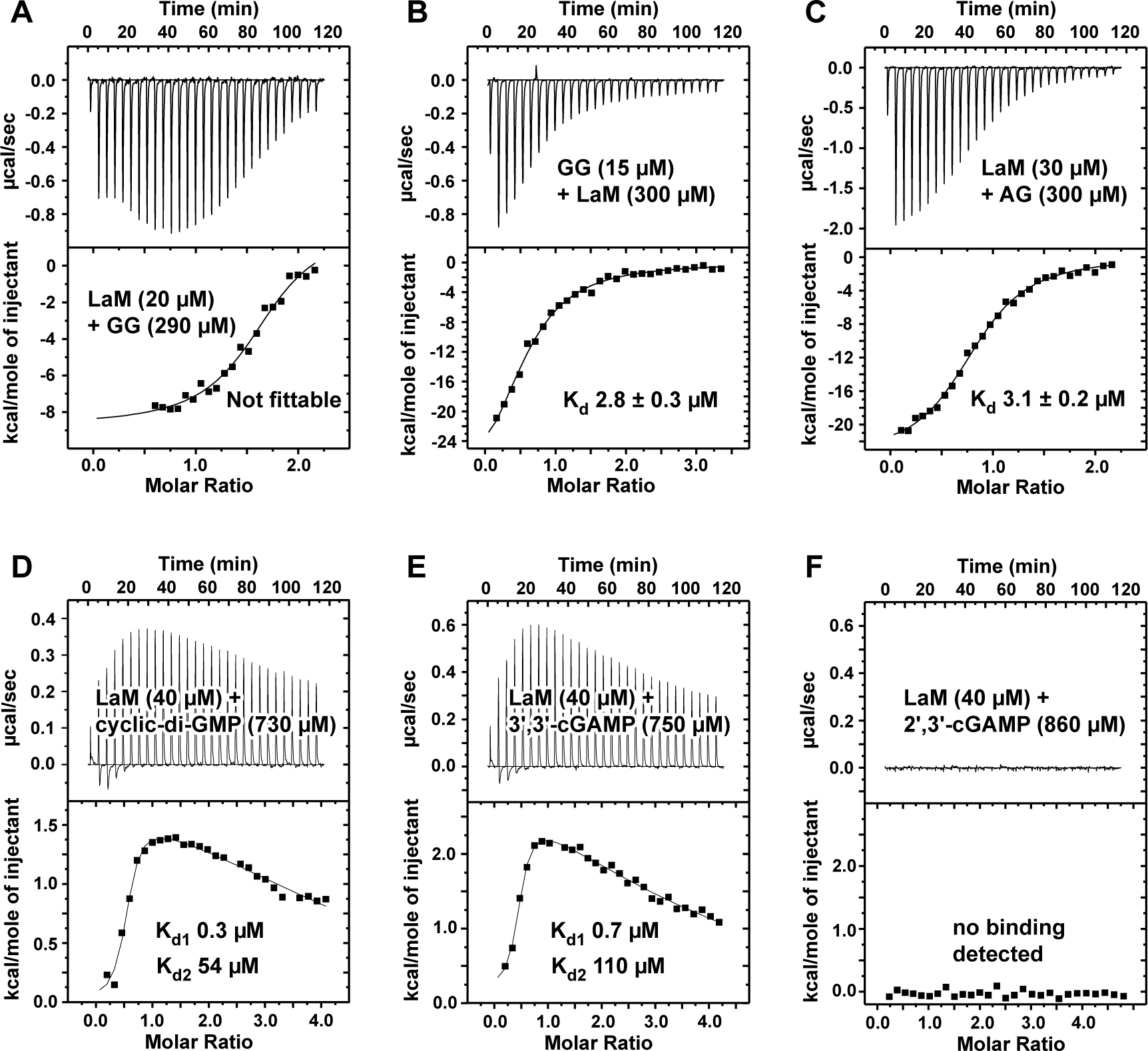
ITC thermograms of dinucleotides binding to LARP1 (323-410). (A) Concentrated 5’-A_P_A- 3’ injected into dilute LaM domain generated a thermogram that could not be fit as simple intermolecular interaction due to competition between guanine quadruplex formation and LaM binding. **(B)** Reversing the components (and concentrations) in the syringe and cell showed the normal behavior for 1-to-1 intermolecular binding. **(C)** Control of concentrated 5’-A_P_G-3’ injected into dilute LARP1 (323-410) showed normal binding. **(D,E)** The second messengers, cyclic-di-GMP and 3’,3’-cGAMP, bound the LaM domain with sub-micromolar affinity. The thermograms were fit using two independent sets of binding reactions. **(F)** The related compound, 2′3′-cGAMP, didn’t show any heat of binding. Either it has no affinity for LaM or the enthalpy of binding is too small to detect.

**Figure 4.**
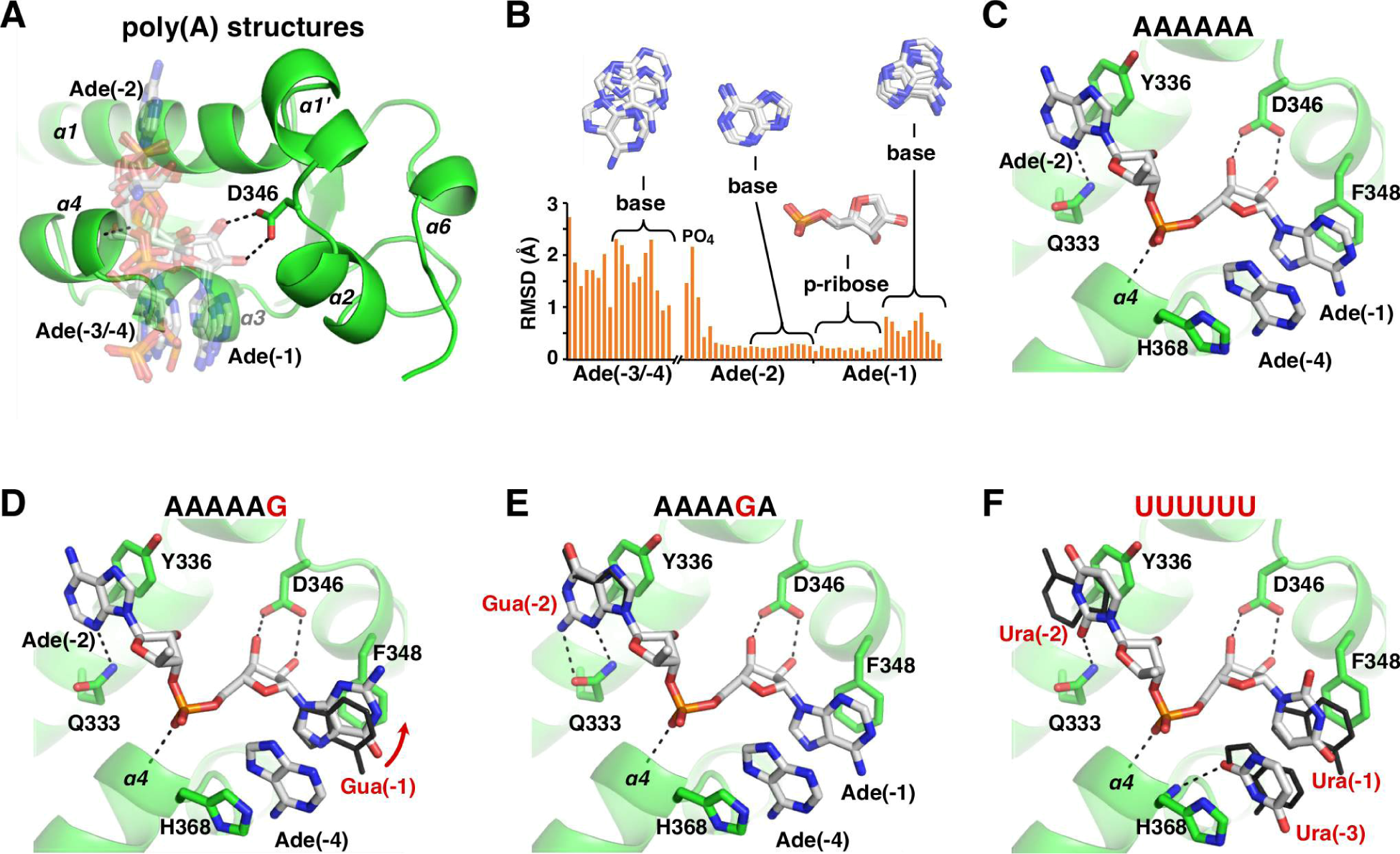
Structures of LARP1 LaM domain in complex with guanosine-containing poly(A) RNAs and poly(U). (A) Overlay of seven structures of poly(A) RNAs: 2×A_3_, A_4_, 2×A_6_, A_3_UA_2_, and A_11_ [7]. Two different stacking conformations are present. In four structures, adenine (-4), numbered from the 3′-end, stacks on adenine (-1), while adenine (-3) does so in the other structures. Specificity for the 3′-terminus is mediated by polar contacts with Asp346. **(B)** Spread of atomic positions in the seven structures. Ribose (-1) and adenine (-2) show the least spread in the overlay. **(C)** Structure of A_6_ bound to LARP1 LaM (PDB 7SOS) [7]. Adenine base (-1) and adenine (-4) stack between Phe348 and His368. Adenine (-2) stacks against Tyr336. (Only selected atoms are shown for clarity.) **(D)** Structure of the complex with A_5_G. Guanine (-1) is shifted up relative to adenine in the A_6_ structure (*black*). The guanine N2 makes an additional hydrogen bond with an ordered sulfate ion in the crystal (*not shown*). **(E)** Structure of the complex with A_4_GA. Guanine base (-2) makes additional hydrogen bond with Gln333, improving the binding affinity. **(F)** Comparison of the complexes with U_6_. The uracil bases overlap with adenines in the A_6_ structure (*black*) but are smaller and contribute less to the binding affinity. Adenine (-4) makes two hydrogen bonds with protein backbone atoms (*not shown*) while uracil (-3) makes only one.

The low micromolar affinity for the dinucleotide GG raised the question if the second messengers in the cyclic GMP-AMP synthase (cGAS)/stimulator of interferon genes (STING) pathway {Ablasser, 2019 #82} could also bind. While the STING agonist, 2′,3′-cGAMP, showed no signal in an ITC titration, both cyclic-di-GMP and 3’,3’-cGAMP appeared to bind with high affinity (Fig. 3D-F). The thermograms showed endothermic binding that required fitting as two sets of independent sites. The initial binding transition was sub-micromolar in affinity with near zero enthalpy change while the second transition was weakly endothermic and very weak affinity. The significance of the complex binding behavior is unclear but the ITC experiments definitely indicate binding of the dinucleotides to the LaM domain and raise the possibility that LARP1 acts downstream of cGAS.

### Structural determinants of base specificity

We turned to X-ray crystallography to understand the molecular basis of LARP1 preference for RNA with guanine substitutions (Table 1, Fig. 3). Co-crystallization trials of LARP1 (323-410) with the RNA hexamers, AAAAAG, AAAAGA, and AAAGAA, yielded crystals diffracting to better than 2 Å (Suppl. Table S2). We also crystallized the protein with RNA with uracil bases, U_6_, and the Rp and Sp- diastereomers of a phosphorothioate derivative of A_6_. The crystals were solved using molecular replacement with the structure of the unliganded LaM domain [7]. All structures showed clear, well-resolved electron density for the protein and 3’-terminal nucleotides (Suppl. Fig. S3).

**Table 1.**
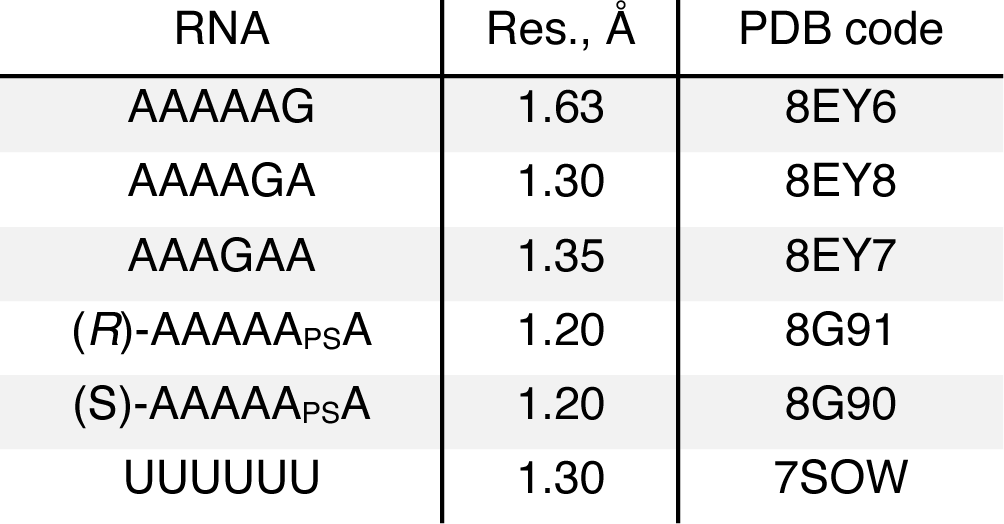
X-ray crystal structures reported.

| RNA | Res., Å | PDB code |
| --- | --- | --- |
| AAAAAG | 1.63 | 8EY6 |
| AAAAGA | 1.30 | 8EY8 |
| AAAGAA | 1.35 | 8EY7 |
| (*R*)-AAAAA<sub>PS</sub>A | 1.20 | 8G91 |
| (*S*)-AAAAA<sub>PS</sub>A | 1.20 | 8G90 |
| UUUUUU | 1.30 | 7SOW |

Not surprisingly, the structures globally resemble the LARP1 LaM domain with bound poly(A) ligands [7]. We previously showed that RNAs can bind in two conformations that differ in the stacking of either the (-3) or (-4) adenine onto adenine (-1). In all the poly(A) structures, the 3’-terminal nucleotides are bound by a pocket formed by alpha helices α1, α2, and α4 (Fig. 3A). Nucleotides (-1) and (-2) are rigidly positioned by hydrogen bonds between the terminal ribose and Asp346, the helical dipole from alpha helix α4, and base stacking with Phe348 and Tyr336. In contrast, the (-3/-4) nucleotide is loosely positioned with large variations between the different structures (Fig. 3B). In the A_6_ structure, the (-4) adenine stacks on His368 while the (-3) nucleotide is completely disordered (Fig. 3C). Comparison of the AAAAAG structure with A_6_ shows the substituted guanine base is shifted to improve the stacking with Phe348 (Fig. 3D). The guanine N2 atom additionally forms an indirect hydrogen bond with the side chain of Arg351. In the crystal, this is formed via a sulfate ion in the crystallization buffer but could be formed via a water molecule observed in the A_4_ structure [7].

The AAAAGA structure clearly explains the 3-fold improvement in binding affinity over A_6_ (Fig. 3E). The introduction of a guanine base adds a hydrogen bond with Gln333. This is consistent with the broadening of the Gln333 side chain NMR resonances observed in the presence of 5’-GMP (Fig. 1A). Previous mutagenesis of Gln333 confirmed that it plays an oversized role in RNA binding with a 20-fold drop in affinity measured for the alanine substitution [7]. mRNA stabilization assays and poly(A) tail- sequencing validated the requirement for Gln333 for LARP1 function in cells. We tested the effects of mutating Gln333 to alanine for binding guanylated poly(A) RNAs. ITC measurements confirmed that loss of the glutamine led to more than a 50-fold loss in binding affinity (Suppl. Table S3).

We also obtained a crystal structure with AAAGAA bound. Unlike the other structures, AAAGAA does not show notable differences relative to the A_6_ structure (Suppl. Fig. S4A). The guanine base does not directly interact with the protein and the conformation of the bound adenine nucleotides is unchanged. Instead, the guanine base (-3) stacks on the 5’-terminal adenine (-6) and makes hydrogen bonds with its own phosphates and those of adenylate (-5). The structure is unusual in that all six nucleosides are visible in the electron density map due to crystal packing and contacts between the RNA 5’-terminal nucleotides (Suppl. Fig S4B).

To understand why the LARP1 LaM binds poly(U) weakly compared to other LARP proteins, we crystalized the domain with U_6_ RNA and determined its structure at 1.30 Å resolution. U_6_ binds LaM with 11 times lower affinity than A_6_ [7]. In the structure, the key binding determinants are conserved although U_6_ adopts the alternative base stacking conformation (-3/-1) with the third nucleotide stacked between the side chain of His368 and uracil (-1). Both (-3/-1) and (-4/-1) stacking configurations likely occur in solution. The crystal structure shows the ribose of nucleotide (-1) is rigidly positioned in the binding pocket formed by Asp346. Uracil bases (-1) and (-2) occupy the same positions as adenines (-1) and (-2) in the poly(A) structure (Fig. 3F). Uracil (-1) stacks on Phe348 and uracil (-2) stacks on Tyr336 with a hydrogen bond to Arg333 [7]. The uracil ring (-3) makes a single hydrogen bond to the protein, between O2 and the backbone amide of His368, compared to two for adenine (-4) in A_6_. Density was observed for the ribose of uridiylate (-4) but it makes no contacts with the protein. Thus, the weaker stacking interactions and fewer protein contacts likely explain the lower affinity of U_6_ compared to A_6_.

### Structural determinants of specificity for phosphoribose

Previous studies have demonstrated the importance of Asp346 and the equivalent aspartic acid in other LARP proteins for RNA binding [7, 24–31]. Conversely, modifications of the 3’ sugar demonstrated the LARP1 LaM is strictly specific for the 3’-terminus. While a deoxyribose sugar is tolerated, blocking of the 3’-terminus with either a 2’-O-methyl or 3’-phosphate group completely prevents binding.^6^ This strict specificity is thought to be a consequence of the tight positioning of phosphoribose in the LaM binding pocket.

To test the importance of the phosphate group, we prepared A_6_ with phosphorothioate modifications at the 3’-terminal nucleotide (-1) and determined the structures of their complexes with LaM at 1.2 Å (Fig. 5). The (*R*_P_)-diastereomer has the sulfur positioned above the binding pocket and reproduces the hydrogen bonding and other contacts seen with A_6_ (Fig. 5A). The phosphate oxygen hydrogen bonds with both the backbone amide of Arg369 and the side chain OH of Tyr337. This positions the 3’-terminal ribose with its hydrophobic face (the face opposite the 2’- and 3’-OH groups) in close contact with the benzyl ring of Phe367. In agreement with the crystal structure, the *R*_P_ isomer binds with essentially the same affinity as A_6_ (290 nM). The (*S*_P_)-diastereomer switches the positions of the oxygen and sulfur in the phosphothioate group and has a dramatic effect on LaM binding. The low electronegativity of sulfur atom makes it a weak hydrogen bonding acceptor [39] and its larger atomic radius displaces the ribose away from Phe367 (Fig. 5C). ITC measurements showed the *S*_P_ isomer binds with 17 µM affinity, confirming the binding selectivity for the *R*_P_ isomer (Fig. 5D).

**Figure 5.**
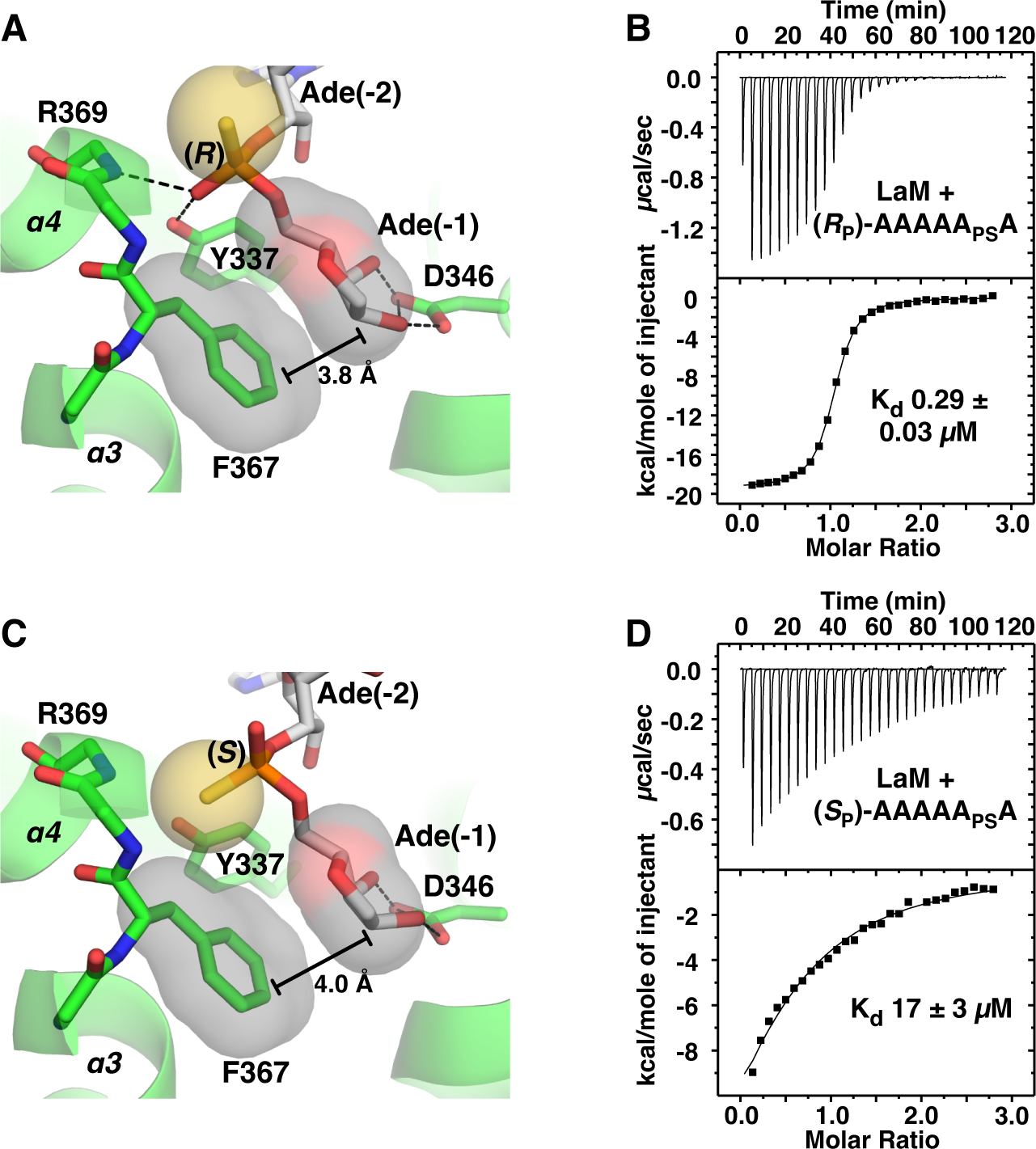
Structure of LARP1 LaM domain with phosphorothioate-containing A_6_ RNA. (**A**) Crystal structure of the complex of the (*R*_P_)-diastereomer 5’-A_P_A_P_A_P_A_P_A_PS_A-3’ shows the sulfur atom (*yellow*) does not interfere with binding. As observed in structures with phosphate, the 3’-ribose is positioned by hydrogen bonds with Asp346 and packing of the sugar aliphatic face against Phe367. The phosphorothioate makes hydrogen bonds with the backbone amide of Arg369 and side chain hydroxyl of Tyr337. (**B**) ITC confirms the (*R*_P_)-phosphorothioate does not affect binding. (**C**) Complex of (*S*_P_)-diastereomer. The sulfur disrupts hydrogen bonds and shifts the ribose away from Phe367. (**D**) ITC shows the (*S*_P_)-phosphorothioate binds with 60-fold lower affinity.

## DISCUSSION

LARP proteins are named after the La autoantigen protein (which was subsequentially renamed LARP3). The family of proteins emerged early in evolution and are present in essentially all eukaryotes [3, 4]. The prototypical LARP gene underwent multiple duplication events to generate specialized isoforms that are conserved across species. It likely contained both LaM and RRM domains since the pair is present in most present day LARP proteins with a few notable exceptions, such as LARP1 [7, 40].

Human LARP1 is large protein (1019 or 1096 amino acids depending on the isoform) with only two independently folding domains. Together, they compose about a quarter of the protein: the LaM domain recognizes RNA 3’-termini and DM15, an alpha-helical HEAT-repeat domain, binds the mRNA 5’-cap and TOP RNA sequences. DM15 domains are unique to LARP1 and LARP1B and serve to negatively regulate TOP mRNA translation in response to mTORC1 inhibition [8, 13, 16, 20, 21, 41]. Unlike the other LARP proteins which contain an RRM domain, the LARP1 LaM domain functions as a stand-alone RNA-binding domain with specificity for poly(A) RNA [7]. While the isolated LaM domain is sufficient for poly(A) binding, the regions adjacent to the La domain also contribute to RNA binding. Al-Ashtal and colleagues [15] observed that LARP1 residues 310–540 bound poly(A) RNA with 40 nM affinity, roughly 6-fold better than the isolated La domain. The LARP1 arginine-glycine (RG) repeat region (residues 250–267 in the 1019-residue isoform) also contributes to RNA binding. Mattijssen et al. [37] showed that a LARP1 fragment including this region exhibited different binding and functional properties. Ogami et al. [34] similarly showed the poly(A) tail lengthening function of LARP1 was less efficient in the absence of residues 249–311. The importance of intrinsically disordered regions was also observed in LARP4 and shown to be primarily responsible for RNA binding [30]. LARP1 also contains a PABP-interaction motif (PAM2) immediately following the LaM domain that serves to recruit PABPC1 and provides increased affinity and specificity for poly(A) mRNA [13, 21, 37]. Uniquely among LARP proteins, LARP1 has the potential to circularize mRNAs through binding of the mRNA 5’-cap by its DM15 domain and the 3’-end by the LaM domain.

Numerous studies have shown that, in addition to regulating TOP mRNA translation, LARP1 protects mRNA with poly(A) tails from degradation [7, 9, 10, 13, 35, 37, 42]. Of particular interest, is the observation that the incorporation of guanine bases (guanylation) into mRNA poly(A) tails by the nucleotidyl transferases TENT4A (PAPD7) and TENT4B (PAPD5) shields them from rapid deadenylation [43, 44]. The accumulation of 3’-terminal guanine nucleotides arises from their slower deadenylation by the CCR4-NOT complex. Our observation of the enhanced affinity of guanine-containing poly(A) sequences suggests the preferential binding of the LaM domain could also contribute to the enrichment of guanine- tailed mRNAs. Comparing lists from Philippe et al. {Philippe, 2020 #60} and Lim et al. {Lim, 2018 #31}, we note that TOP mRNAs were markedly over-represented in the list of guanylated mRNAs. Two-thirds of TOP mRNAs (63 of 97) were in the set of 390 mRNAs observed to be guanylated in 5 or more reads. Conversely, 16% (62 of 390) of guanylated mRNAs were TOP mRNAs. RPSA (small ribosomal subunit protein uS2) was the most frequently guanylated TOP mRNA with close to one in twenty sequences terminating with guanine or a penultimate guanine nucleotide. Notably, LARP1 was shown to interact with TENT4A and TENT4B to induce post-transcriptional polyadenylation of TOP mRNAs [34] and act with PABPC1 as a barricade to deadenylation by CCR4-NOT [35].

Surprisingly, within the list of guanylated mRNAs, TOP mRNAs showed less frequent guanylation than the average. mRNAs for secreted and endoplasmic reticulum proteins ended in guanine more frequently than the ribosomal proteins and translation factors [44]. This could reflect dynamic guanylation of TOP mRNAs downstream of mTOR signaling. Alternatively, it may reflect mTOR-independent functions of LARP1. TOP mRNAs account for less than 4% of mRNAs bound by LARP1 in cancer cells [14, 18, 19].

### LaM domain binding specificity

The observation of 2 to 3-fold increased affinity for single guanine-containing RNAs led to the expectation that multiple guanine bases would further increase the affinity (Fig. 2B). Instead, there is no improvement and, in the case of the ligands AAAAGA (80 nM) and AAAAGG (140 nM), the second guanine leads to a loss of affinity. Similar behavior was observed with the dinucleotides AG and GG which both bind with ∼3 µM affinity (Fig. 3). The biophysical basis for the negative cooperativity of the guanines is unclear. In general, relatively small (two-fold) differences in binding affinities are difficult to rationalize based on static crystal structures. The crystal structure of AAAGAA bound to the LaM does little to explain the 1.8-fold improvement in affinity. While comparison of the A_6_ and AAAGAA complexes shows a small shift in the position of the (-3) adenine ring (Suppl. Fig. S4A), RNA bases at that position are highly variable (Fig. 3B) and the observed shift is unlikely to be fully responsible for the improved affinity. As an alternative explanation, in solution, the guanine base may bind in the (-3/-1) conformation with better stacking and hydrogen bonding interactions than the adenine base. In our study of poly(A) structures, we observed considerable variability in the stacking configurations (-3/-1 versus -4/-1). In one case the same RNA different crystals bound in two different configurations [7]. Multiple conformations are likely to exist in solution. Finally, comparison of the affinity different RNA sequences assumes that the free energies of the unbound RNAs are the same. This is clearly not the case for all guanine RNAs (G_2_ and G_6_), which can form guanine quadruplexes in competition with LaM binding. Uniquely, in the AAAGAA complex, the 5’- nucleotides make intra-RNA contacts and are visible in the electron density map (Suppl. Fig. S4B).

The crystal structure of U_6_ complex does a better job explaining the lower affinity which results from fewer hydrogen bonds and smaller stacking interfaces. U_6_ bound to the LaM domain in a -3/-1 stacking conformation which is consistent with the structure reported for 3’-end poly(U) sequences interacting with LARP3 [26]. In LARP3, the La-module (the combination of LaM and RRM domains) binds the UUU 3ʹ termini of nascent RNA polymerase III transcripts, most of which are precursors to the tRNAs, protecting them from untimely digestion by 3’ exonucleases while also assisting their folding [25, 45, 46]. The U_6_ structure underlines the plasticity in RNA recognition by the LARP1 LaM domain, which lacks the intermolecular contacts and binding specificity provided by the RRM domain found in other LARPs.

Ultimately, the key function of LaM domains appears to be the recognition of the RNA 3ʹ-terminus. This is dramatically illustrated by the comparison of the NMR chemical shift changes from 3ʹ-AMP and 5ʹ- AMP (Fig. 1A). Moving the phosphate to the 3ʹ carbon completely eliminated binding adjacent to I364. The effect is much larger than base modifications. While the role of Asp346 in recognizing the 3ʹ-hydroxyl was known, the importance of contacts between the (-1) phosphate and helix α4 had not been tested previously. We took advantage of chiral phosphorothioate derivatives to demonstrate the helix dipole and hydrogen bonds to backbone amides are major contributors to binding affinity. The strong chiral selectivity we observed opens the door to controlling the stability of synthetic RNAs through stereoselective phosphorothioate derivatives [47, 48].

## MATERIALS AND METHODS

### Expression and purification of proteins

The plasmid expressing human LARP1 LaM domain (residues 323-410) and its purification as a His-tag fusion protein from *E. coli* cells were described previously [7]. Amino acids are numbered according to the shorter 1019-residue isoform.

### RNA oligonucleotides

RNA oligonucleotides A_6_ (5’-A_P_A_P_A_P_A_P_A_P_A-3’), A_5_G (5’-A_P_A_P_A_P_A_P_A_P_G-3’), A_4_GA (5’-A_P_A_P_A_P_A_P_G_P_A-3’), and A_3_GA_2_ (5’-A_P_A_P_A_P_G_P_A_P_A-3’) were synthesized on 2 × 2 µmol scale using an ABI 3400 DNA synthesizer employing the β-cyanoethylphosphoramidite method with long chain alkylamine controlled pore glass (LCAA-CPG, 500 Å) as the solid support functionalized with a protected 5’-O- dimethoxytrityl ribonucleoside. Hexamers containing a phosphorothioate (thiophosphate) internucleotide linkage between the terminal 3’ adenylate residues (5’-A_P_A_P_A_P_A_P_A_PS_A-3’) were prepared using the sulfurization reagent 3-phenyl 1,2,4-dithiazoline-5-one (PolyOrg Inc.) at a concentration of 0.1 M in acetonitrile and 2 equivalents of pyridine for a total exposure of 2 min after the coupling of the first β- cyanoethylphosphoramidite in the synthesis cycle. After this step, solid-phase synthesis proceeded according to previously published procedures [49]. Oligonucleotides were cleaved from the solid support, deprotected, purified by either preparatory denaturing PAGE or ion-exchange HPLC (for the separation of the (*R*_P_)- and (*S*_P_)-diastereomers of 5’-A_P_A_P_A_P_A_P_A_PS_A-3’), and then desalted using C-18 SEP PAK cartridges. The dinucleotides, 5’-A_P_G-3’ and 5’-G_P_G-3’, were obtained from Jena Bioscience (Thuringia, Germany). U_6_ (5’-U_P_U_P_U_P_U_P_U_P_U-3’) was obtained from Integrated DNA Technologies (Coralville, USA) and used without additional purification. cyclic-di-GMP (CAS no. 61093-23-0), 3’,3’-cGAMP (CAS no. 849214-04-6), and 2’,3’-cGAMP (CAS no. 1441190-66-4) were obtained from Sigma-Aldrich and used without further purification.

### NMR spectroscopy

NMR samples were exchanged into 10 mM MES pH 6.3, 100 mM NaCl, 1 mM TCEP. For NMR titrations, nucleotides were added to 0.1 mM ^15^N-labeled LARP1 (323-410) to final molar ratios of 5:1. All NMR experiments were performed at 25 °C using a Bruker 600 MHz spectrometer. NMR spectra were processed using NMRPipe [50] and analyzed with SPARKY [51]. Binding affinities were measured by fitting the observed chemical shift changes accounting for bound ligand with a stoichiometry of two nucleotide binding sites.

### Isothermal titration calorimetry

ITC experiments were performed on the VP-ITC titration calorimeter (Malvern Instruments Ltd) except where noted. The syringe was loaded with 300 µM RNA sample, while the sample cell contained 20-30 µM LARP1 (323-410) sample. All experiments were carried out at 20 °C with 29 injections of 10 µl. Results were analyzed using ORIGIN software (MicroCal) and fitted to a binding model with a single set of identical sites, except where noted.

### Crystallization

Initial crystallization conditions were identified utilizing hanging drop vapor diffusion with the Classics II and Nucleix screens (QIAGEN). The best LARP1 LaM domain/RNA complex crystals were obtained by equilibrating a 0.6 µL drop of the LaM domain (residues 323-410) with oligonucleotide in a 1:1.1 molar ratio (10 mg/mL of protein) in buffer (10 mM MES pH 6.3, 100 mM NaCl), mixed with 0.6 µL of reservoir solution containing [0.1 M Bis-Tris pH 6.5, 2 M ammonium sulfate] for A_5_G, [0.2 M ammonium acetate, 0.1 M Bis-Tris pH 6.5, 25% (w/v) PEG 3350] for A_4_GA, [0.056 M sodium phosphate, 1.344 M potassium phosphate] for A_3_GA_2_, [0.2 M ammonium acetate, 0.1 M Bis-Tris pH 5.5, 25% (w/v) PEG 3350] for U_6_, [0.1 M ammonium acetate, 0.1 M Bis-Tris pH 5.5, 17% (w/v) PEG 10000] for (*R*_P_)- A_P_A_P_A_P_A_P_A_PS_A, [0.1 M Bis-Tris pH 6.5, 28% (w/v) PEG 2000 MME] for (*S*_P_)-A_P_A_P_A_P_A_P_A_PS_A. Crystals grew in 3-14 days at 20 °C. For data collection, crystals were cryo-protected by soaking in the reservoir solution supplemented with 30% (v/v) ethylene glycol for conditions PEG or otherwise with 25% glycerol.

### Structure solution and refinement

Diffraction data from single crystals of LARP1 LaM domain and its RNA complexes were collected at the Canadian Light Source (CLS) and Cornell High-Energy Synchrotron Source (CHESS). Data processing and scaling were performed with HKL2000 [52]. The initial phases for the complex structures were determined by molecular replacement with Phaser [53], using the coordinates of the LARP1 LaM domain (PDB entry 7SOO) [7]. The starting protein model was then completed and adjusted with the program Coot [54] and improved by several cycles of refinement, using the program phenix.refine [55] and model refitting. The resulting electron density maps revealed clear density for each RNA oligonucleotide, which were manually built with the program Coot [54]. At the latest stage of refinement, we also applied the translation-libration-screw (TLS) option [56]. The final models have all residues in the allowed regions of Ramachandran plot. Data collection and refinement statistics are given in Suppl. Table S2.

## ACKNOWLEDGEMENTS

Crystallographic data were acquired at the Canadian Light Source (CLS) and the Cornell High-Energy Synchrotron Source (CHESS) Macromolecular Diffraction facility. CLS is supported by the Canada Foundation for Innovation, the Natural Sciences and Engineering Research Council, the National Research Council, the Canadian Institutes of Health Research, and the Government of Saskatchewan. CHESS is supported by National Science Foundation award DMR-0225180 and National Institutes of Health/NCRR award RR-01646. The study has been supported by funding from Canadian Institutes of Health Research grant FDN 159903 to K.G.

## DATA AVAILABILITY

The authors confirm that the crystallographic data are available from the Protein Data Bank (PDB) under accession codes 8EY6 (A_5_G), 8EY8 (A_4_GA), 8EY7 (A_3_GA_2_), 8G91 (A_6_ with *R*_P_-phosphorothioate diastereomer), 8G90 (A_6_ with *S*_P_- phosphorothioate diastereomer), and 7SOW (U_6_). Other data supporting the findings of this study are available within the article and its supplementary materials.

