## Supplemental Tables and Figures for "Enhanced binding of guanylated poly(A) RNA by the LaM domain of LARP1"

**Table of Contents**

Table S1. ITC Data Collection and Analysis for LARP1 323-410 page 2

Table S2. X-ray Data Collection and Refinement Statistics page 3

Table S3. ITC Data Collection and Analysis for LARP1 323-410 Q333A page 5

Figure S1. $^{15}N$-$^{1}H$ correlation spectra of $^{15}N$-labeled LARP1 (323-410) page 6

Figure S2. ITC thermograms of binding between LARP1 LaM and RNAs page 7

Figure S3. Representative electron density page 8

Figure S4. Crystal structure of LARP1 LaM with $A_3GA_2$ RNA oligonucleotide bound page 9

Table S1. ITC Data Collection and Analysis for LARP1 323-410

| RNA | $K_d$, nM | Affinity relative to $A_6$ | ΔH, kcal $mol^{-1}$ | ΔS, cal $mol^{-1}$ $K^{-1}$ | Conc., µM |
|---|---|---|---|---|---|
| **AAAAAA** | 250 ± 15 | 1 | -19.9 | -37.6 | LaM 30 (cell), RNA 300 (syringe) |
| **AAAAAG** | 130 ± 20 | 0.52 | -27.5 | -62.3 | LaM 20 (cell), RNA 300 (syringe) |
| **AAAAGA** | 80 ± 10 | 0.32 | -29.5 | -68.0 | LaM 20 (cell), RNA 300 (syringe) |
| **AAAGAA** | 140 ± 20 | 0.56 | -27.7 | -63.2 | LaM 20 (cell), RNA 300 (syringe) |
| **AAGAAA** | 140 ± 10 | 0.56 | -29.3 | -68.8 | LaM 30 (cell), RNA 300 (syringe) |
| **AAAAGG** | 140 ± 10 | 0.56 | -27.4 | -62.3 | LaM 20 (cell), RNA 300 (syringe) |
| **AAAGAG** | 120 ± 10 | 0.48 | -31.2 | -74.7 | LaM 20 (cell), RNA 300 (syringe) |
| **AAGAGA** | 70 ± 10 | 0.28 | -38.3 | -97.9 | LaM 20 (cell), RNA 300 (syringe) |
| **(*R*$_P$)-AAAAA$_{PS}$A** | 290 ± 30 | 1.16 | -19.4 | -36.2 | LaM 25 (cell), RNA 300 (syringe) |
| **(*S*$_P$)-AAAAA$_{PS}$A** | 17000 ± 2700 | 68 | -23.1 | -57.1 | LaM 25 (cell), RNA 300 (syringe) |
| **UUUUUU** | 1800 ± 100 | 7.2 | -21.4 | -46.6 | LaM 30 (cell), RNA 300 (syringe) |
| **GG** | not fitted | n/a | n/a | n/a | LaM 20 (cell), RNA 290 (syringe) |
| **GG** | 2800 ± 300 | 11.2 | -34.1 | -91.1 | RNA 15 (cell), LaM 300 (syringe) |
| **AG** | 3100 ± 200 | 12.4 | -23.9 | -56.2 | LaM 30 (cell), RNA 300 (syringe) |
| **cyclic-di-GMP (site 1)**<br>**(site 2)** | 300 ± 300<br>>10000 | 1.2<br>220 | 0<br>2.0 | 29.8<br>26.5 | LaM 40 (cell), ligand 730 (syringe) |
| **3',3'-cGAMP (site 1)**<br>**(site 2)** | 700 ± 300<br>>10000 | 2.8<br>440 | 0<br>4.8 | 28.3<br>34.4 | LaM 40 (cell), ligand 750 (syringe) |
| **2',3'-cGAMP** | not fitted | n/a | n/a | n/a | LaM 40 (cell), ligand 860 (syringe) |

Table S2. Data Collection and Refinement Statistics

| **Data collection** | LaM-$A_5G$ | LaM-$A_4GA$ | LaM-$A_3GA_2$ |
|---|---|---|---|
| PDB code | 8EY6 | 8EY8 | 8EY7 |
| Space group | $P2_12_12_1$ | $P2_12_12_1$ | $P2_12_12_1$ |
| Cell dimensions | | | |
| *a*, *b*, *c* (Å) | 36.52, 46.19, 58.90 | 36.68, 45.88, 58.86 | 36.91, 45.34, 59.41 |
| Resolution (Å) | 50-1.63 (1.66-1.63)[1] | 50-1.30 (1.32-1.30) | 50-1.35 (1.37-1.35) |
| $R_{sym}$ | 0.091 (0.537) | 0.106 (0.427) | 0.109 (0.498) |
| $I / \sigma I$ | 16.6 (1.5) | 22.1 (2.2) | 17.7 (1.7) |
| Completeness (%) | 97.2 (96.3) | 95.5 (65.5) | 93.3 (57.6) |
| Redundancy | 6.0 (4.8) | 6.4 (2.8) | 7.1 (5.0) |
| CC1/2[2] | 0.782 | 0.932 | 0.967 |
| **Refinement** | | | |
| Resolution (Å) | 36.3 - 1.63 | 36.2 - 1.30 | 28.6 - 1.35 |
| No. reflections | 12601 | 23885 | 20855 |
| $R_{work}$ / $R_{free}$ | 0.208/0.242 | 0.183/0.192 | 0.186/0.225 |
| No. atoms | | | |
| Protein | 723 | 746 | 784 |
| RNA | 77 | 77 | 130 |
| Water | 33 | 62 | 81 |
| *B*-factors | | | |
| Protein | 30.2 | 22.0 | 16.9 |
| RNA | 51.6 | 35.4 | 21.7 |
| Water | 35.8 | 29.9 | 24.4 |
| R.m.s deviations | | | |
| Bond lengths (Å) | 0.007 | 0.014 | 0.013 |
| Bond angles (°) | 0.98 | 1.59 | 1.46 |
| Ramachandran statistics (%) | | | |
| Most favored regions | 97.7 | 97.7 | 97.8 |
| Additional allowed regions | 2.3 | 2.3 | 2.2 |
| Disallowed regions | 0.0 | 0.0 | 0.0 |

[1]Highest resolution shell is shown in parentheses.
[2]CC1/2 in highest resolution shell.

Table S2. Data Collection and Refinement Statistics (continued)

| **Data collection** | LaM-$A_5$($R_P$)A[3] | LaM-$A_5$($S_P$)A[3] | LaM-$U_6$ |
|---|---|---|---|
| PDB code | 8G91 | 8G90 | 7SOW |
| Space group | $P4_32_12$ | $P2_12_12_1$ | $P2_12_12_1$ |
| Cell dimensions | | | |
| $a$, $b$, $c$ (Å) | 53.78, 53.78, 90.45 | 36.95, 46.65, 59.28 | 36.80, 46.89, 57.90 |
| Resolution (Å) | 50-1.20 (1.22-1.20) | 50-1.20 (1.22-1.20) | 50-1.30 (1.32-1.30) |
| $R_{sym}$ | 0.104 (1.37) | 0.081 (0.445) | 0.060 (0.431) |
| $I / \sigma I$ | 42.4 (1.2) | 35.6 (2.8) | 24.7 (1.6) |
| Completeness (%) | 99.4 (96.6) | 95.2 (67.7) | 99.1 (86.7) |
| Redundancy | 22.1 (10.3) | 8.6 (2.9) | 7.3 (3.7) |
| CC1/2 | 0.592 | 0.825 | 0.866 |
| **Refinement** | | | |
| Resolution (Å) | 35.1 - 1.20 | 26.0 - 1.20 | 36.4 - 1.30 |
| No. reflections | 41809 | 31018 | 24922 |
| $R_{work}$ / $R_{free}$ | 0.178/0.189 | 0.192/0.209 | 0.182/0.192 |
| No. atoms | | | |
| Protein | 749 | 737 | 797 |
| RNA | 75 | 75 | 69 |
| Water | 106 | 115 | 66 |
| $B$-factors | | | |
| Protein | 17.6 | 17.0 | 20.8 |
| RNA | 33.3 | 29.8 | 29.8 |
| Water | 26.4 | 29.2 | 27.6 |
| R.m.s deviations | | | |
| Bond lengths (Å) | 0.006 | 0.004 | 0.014 |
| Bond angles (°) | 1.17 | 0.94 | 1.49 |
| Ramachandran statistics (%) | | | |
| Most favored regions | 97.7 | 97.7 | 96.8 |
| Additional allowed regions | 2.3 | 2.3 | 3.2 |
| Disallowed regions | 0.0 | 0.0 | 0.0 |

[3]Phosphorothioate RNA

Table S3. ITC Data Collection and Analysis for LARP1 323-410 Q333A

| RNA | $K_d$, nM | Affinity relative to wild-type LARP1 | ΔH, kcal mol$^{-1}$ | ΔS, cal mol$^{-1}$ K$^{-1}$ | Conc., µM |
|---|---|---|---|---|---|
| **AAAAAG** | 19000 ± 2000 | 146 | -27.8 | -73.2 | LaM 30 (cell), RNA 300 (syringe)[1] |
| **AAAAGA** | 10000 ± 6000 | 125 | -10.2 | -11.9 | LaM 30 (cell), RNA 300 (syringe)[1] |
| **UUUUUU** | 100000 ± 30000 | 56 | -11.0 | -19.2 | LaM 30 (cell), RNA 600 (syringe) |

[1]Data acquired on iTC200 (Malvern Instruments Ltd).

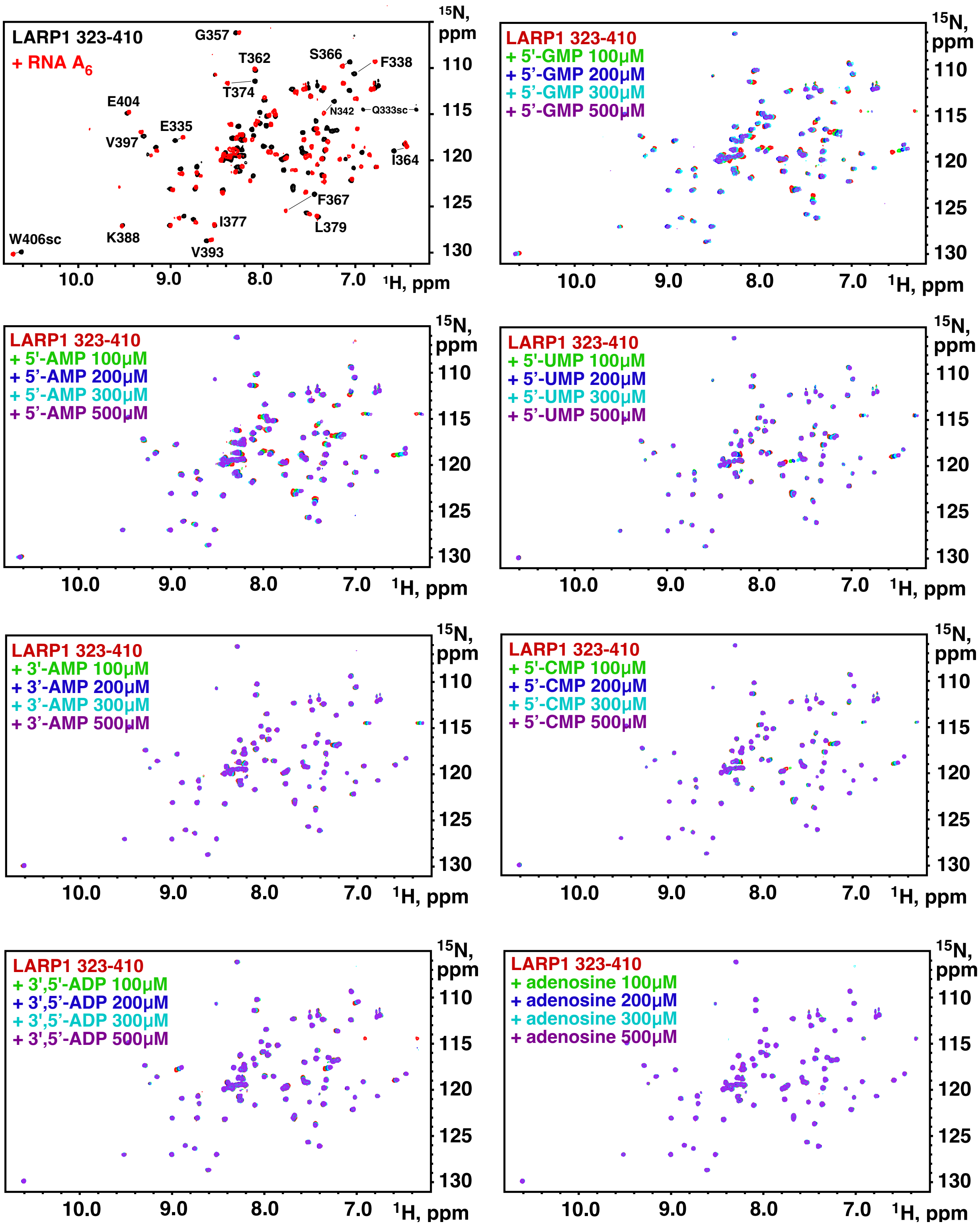


**Supplementary Figure S1**. $^{15}$N-$^{1}$H correlation spectra of $^{15}$N-labeled LARP1 (323-410) in the presence of RNA and single nucleotides. Peak assignments shown for $A_6$ are from reference [1]. sc = side chain.

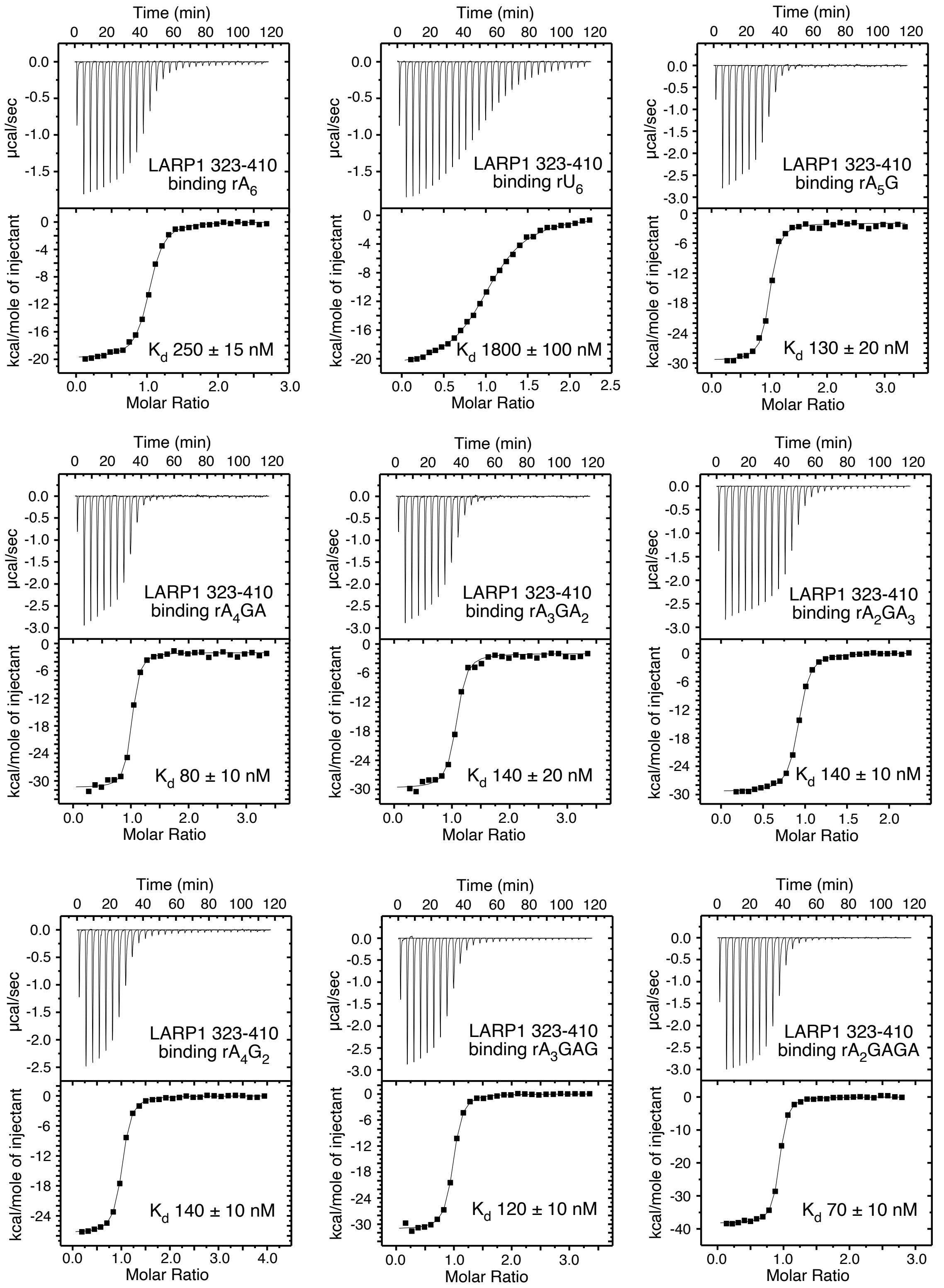


**Figure S2**. ITC thermograms of binding between LARP1 LaM and RNAs.

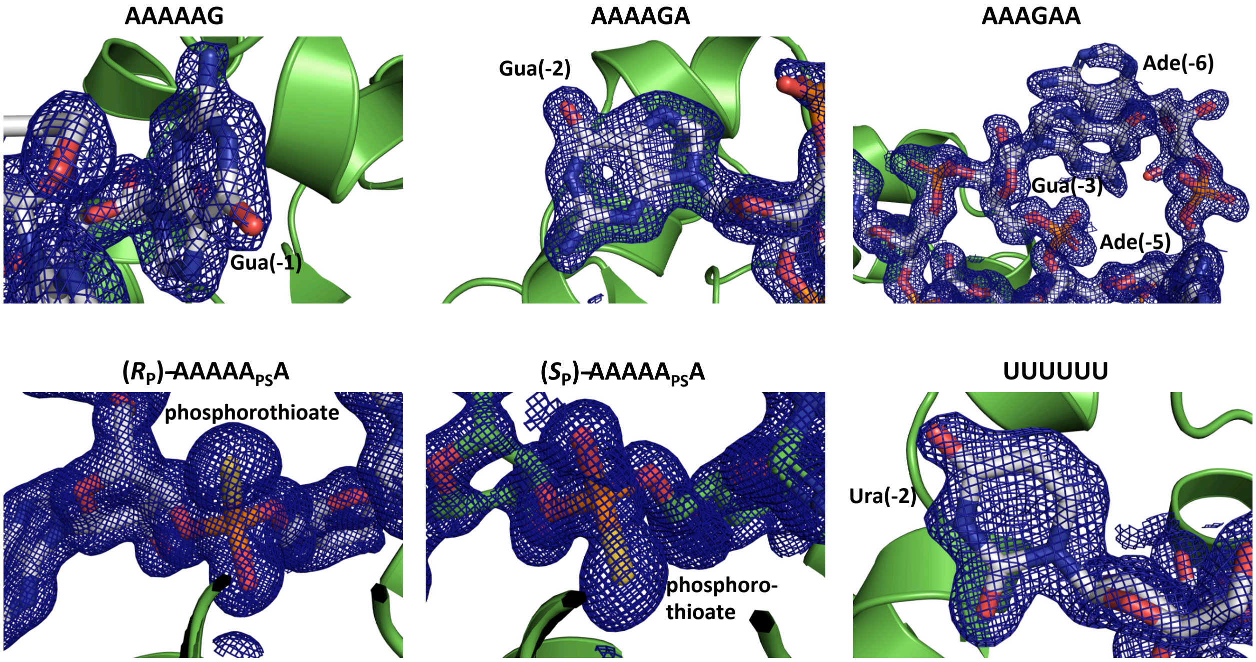


**Figure S3**. Representative electron density. Density is contoured at 1σ from RNA 2Fo-Fc omit maps.

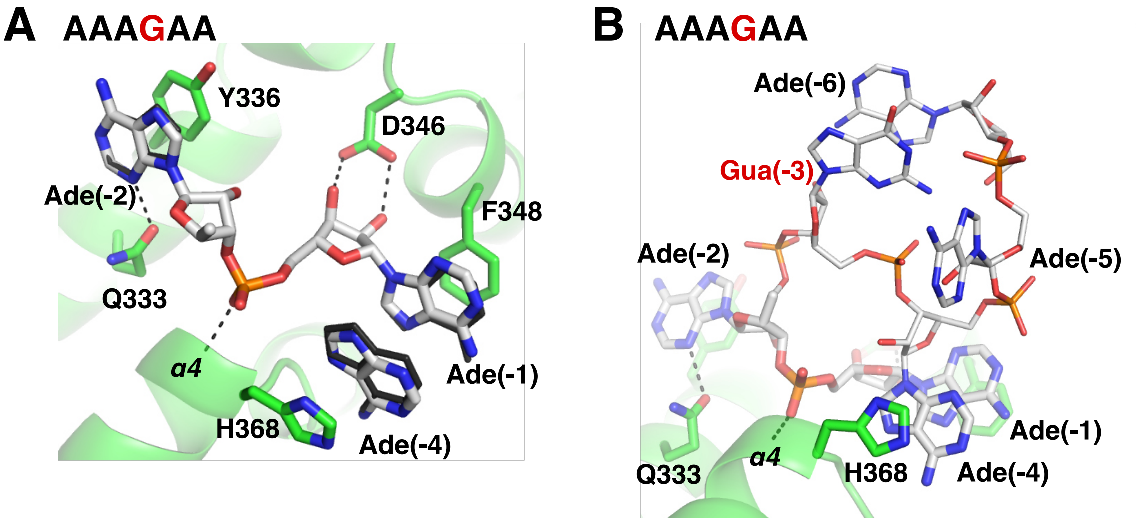


**Figure S4**. Crystal structure of LARP1 LaM with $A_3GA_2$ RNA oligonucleotide bound. **(A)** Comparison of the complex with $A_6$ (*black*). The two structures overlap tightly with only a small shift in the adenine ring at position (-4). (For clarity, only selected atoms are shown.) **(B)** Full model of $A_3GA_2$. The guanine base at position (-3) does not contact the protein but rather stacks against the base of adenylate (-6).
